# gFET-based aptasensors technology allows sensitive and specific quantification of the ESKAPE pathogens

**DOI:** 10.64898/2025.12.12.693864

**Authors:** Shengnan He, Daniel Gruber, Jan-Christoph Walter, Runliu Li, Roger Hasler, Wolfgang Knoll, Christoph Kleber, Dennis Kubiczek, Verena Vogel, Barbara Spellerberg, Ann-Kathrin Kissmann, Frank Rosenau

**Affiliations:** Institute of Pharmaceutical Biotechnology, Ulm University, Albert-Einstein-Allee 11, 89081 Ulm, Germany; Faculty of Medicine and Dentistry, Danube Private University, Steiner Landstraße 124, 3500 Krems an der Donau, Austria; ecoSPECS GmbH, Hermann-Volz-Straße 56, 88400 Biberach an der Riß, Germany; Institute of Medical Microbiology and Hygiene, University Clinic of Ulm, Albert-Einstein-Allee 11, 89081 Ulm, Germany

## Abstract

The rapid rise of antimicrobial resistance among the ESKAPE pathogens, *Enterococcus faecium, Staphylococcus aureus, Klebsiella pneum*oniae, *Acinetobacter baumannii, Pseudomonas aeruginosa* and *Enterobacter* spp., requires diagnostic technologies capable of fast, simplified and sensitive microbial detection. Conventional culture-based diagnostics remain too time-consuming to guide early therapeutic interventions. Here, we present a graphene field-effect transistor (gFET) aptasensor platform enabling rapid, label-free and highly sensitive quantification of all six ESKAPE pathogens. Each gFET device was functionalized with a specific DNA aptamer selected from literature sources and immobilized via a mixed pyrene-based linker strategy on reduced graphene oxide (rGO). Exposure of the functionalized sensors to logarithmically diluted bacterial suspensions (10-10^5^ CFU mL^−^) produced characteristic and concentration-dependent shifts in source–drain current (ΔI_DS_). For all pathogen-specific aptamers, ΔI_DS_ correlated linearly with bacterial load (R^2^ = 0.90-1.00), while non-target bacteria generated only low-level, unspecific fluctuations. Limits of detection ranged from 10 to 1000 bacterial cells depending on the aptamer. Together, these results demonstrate that aptamer-functionalized rGO-FETs provide a robust, scalable and highly specific electronic biosensing architecture capable of distinguishing clinically relevant multidrug-resistant pathogens with excellent analytical performance.

**Author summary:** Antimicrobial-resistant bacteria pose a growing threat to global health, especially the so-called ESKAPE pathogens, which frequently cause hospital-acquired infections and are increasingly difficult to treat. Current diagnostic methods can take several days, delaying the start of effective therapy. In our work, we developed a fast and highly sensitive biosensor that uses electrically conductive GO and short DNA molecules, called aptamers, to recognize specific bacteria. When a pathogen binds to its matching aptamer on the sensor surface, the electrical signal of the graphene changes in a measurable way. We tested ESKAPE species and showed that all of the investigated aptamers detect only their intended bacterial targets, even at very low concentrations. Importantly, the sensors respond within minutes and do not require any labelling or complex sample preparation. Our technology demonstrates how graphene-based aptasensors can support rapid and accurate detection of dangerous bacterial pathogens and could ultimately help clinicians make faster decisions in treating infections.

## Introduction

The alarming increase in antimicrobial resistances (AMRs) represents an urgent and pervasive problem to global public health. Rice et al. introduced the term ESKAPE to designate a group of bacterial pathogens of particular concern, which include *Enterococcus faecium, Staphylococcus aureus, Klebsiella pneumoniae, Acinetobacter baumannii, Pseudomonas aeruginosa* and *Enterobacter*-spp. that are in special focus of multi-resistances (1). These pathogens are notorious for evading (“escaping”) the effects of most commonly used antibiotics and together account for a significant proportion of hospital-acquired infections in healthcare facilities across nearly every continent, with countries such as USA (2), Mexico (3), Brazil (4), Germany (5), Italy (6), Romania (7)(8), Zambia (9), Iran (10), China (11) and Australia (12) serving as illustrative examples of hospitals where these organisms are frequently isolated. Clinically, ESKAPE pathogens are associated with a broad spectrum of diseases. *Enterococcus faecium*, frequently exhibiting vancomycin resistance (VRE), is commonly implicated in bloodstream infections, urinary tract infections and endocarditis (13)(14)(15). *Staphylococcus aureus*, including methicillin-resistant strains (MRSA), is a major cause of skin and soft tissue infections, pneumonia, sepsis and device-associated infections (16)(17)(18)(19). In the case of *Klebsiella pneumoniae*, pneumonia and urinary tract infections are prominent, with many isolates producing extended-spectrum β-lactamases (ESBLs) or carbapenemases (20)(21)(22). Ventilator-associated pneumonia, wound infections and bloodstream infections are typical manifestations of *Acinetobacter baumannii*, with multidrug-resistant strains posing particular challenges (23)(24). Amongst all healthcare-associated infection *Pseudomonas aeruginosa* has a prevalence of 7.1% - 7.3% frequently causing pneumonia and cystic fibrosis (25)(26)(27). The ESKAPE species *Enterobacter* spp. has according to intensive care units a prevalence of 11.1% in respiratory tract isolates and 10.3% of bacterial pathogens recovered from surgical wounds. With a frequency of 6.1% in urinary tract and 5.3% in blood cultures *Enterobacter* spp. have a high prevalence in many different habitats (28). Due to this clinical relevance and their alarming resistance profiles, the ESKAPE pathogens are classified by the World Health Organization (WHO) (29) and the Center for Disease Control (CDC) (30) as high-priority threats requiring new diagnostic and therapeutic strategies. Conventional detection methods such as bacterial culture and biochemical assays remain the gold standard for species identification, but their slow turnaround time of 24-72 hours often delays the initiation of effective therapy. To overcome these limitations, we introduced an aptamer-based rGO-FET biosensor platform that enables rapid and highly sensitive detection of microbial pathogens (31)(32)(33)(34)(35). In this system, aptamers are immobilized on the rGO and serve as the molecular recognition element. Aptamers are short, single-stranded DNA or RNA oligonucleotides capable of folding into defined three-dimensional structures that bind their targets with antibody-like affinity and specificity (36)(37)(38). The rGO-FET biochip translates these binding events into measurable electronic signals, where a rGO channel connects source and drain electrodes, while an applied gate voltage modulates its conductivity. When target molecules bind to the aptamers, the resulting change in local charge distribution perturbs the carrier density in the channel, causing a measurable shift in the source-drain current. To implement this principle in a robust sensing architecture, the gFET devices were functionalized with aptamers through a defined surface-chemistry cascade (Fig 1A). Following functionalization, the biochips are mounted into a microfluidic flow chamber (Fig 1B), enabling controlled rinsing, activation and kinetically defined exposure to the analyte. The resulting binding events are monitored by recording the transfer characteristics of the device (I_DS_V_G_ curves). As depicted in Fig 1C, each functionalization step and subsequent target interaction induces characteristic shifts in the electronic response of the graphene channel, which represent the basis for quantitative biosensing in the following experiments.

**Fig 1:**
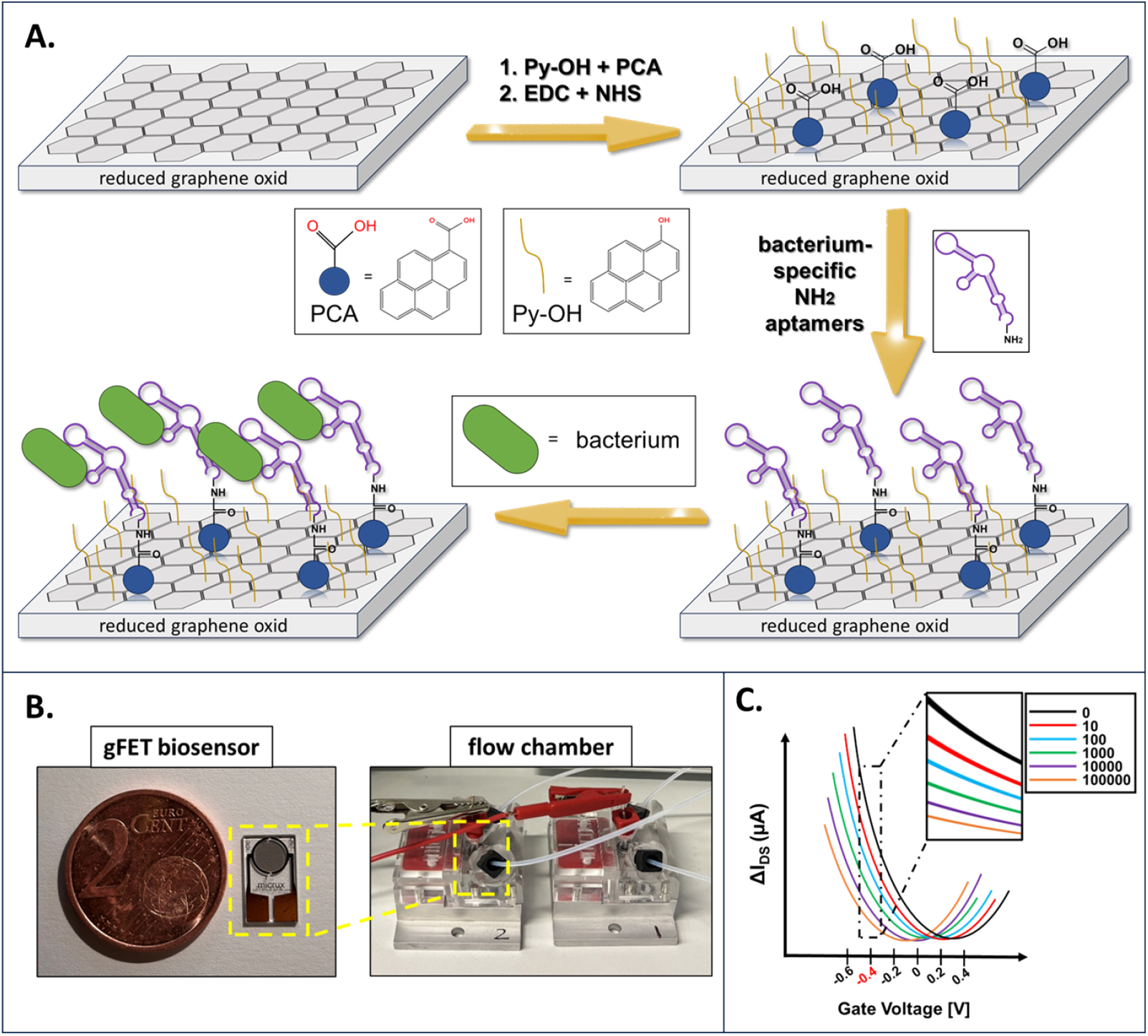
Surface modification, device assembly and signal electrical response of the Aptamer-Based rGO gFET biosensor. (A) Stepwise functionalization of the rGO channel on the FET. Bare rGO is first modified with a mixed Hydroxypyrene (Py-OH)/1-pyridine carboxylic acid (PCA) layer, in which Py-OH passivates the surface and PCA provides carboxyl groups used to covalently couple amino-terminated aptamers, yielding a dense aptamer layer on the rGO surface. The immobilized aptamers subsequently bind their specific bacterial targets, which leads to a change in the local charge environment above the graphene channel. (B) Photograph of the fabricated gFET biosensor and its placement inside a microfluidic flow chamber. (C) Model I_DS_V_G_ transfer curves based on fictive and idealized measurements of the gFET recorded at different stages of the experiment: after initial conditioning of the bare rGO device and after incubation with increasing concentrations of the bacterial target. Each step induces a systematic shift in the transfer curve. For subsequent quantitative analysis, a fixed gate voltage of -0.5 V was chosen (vertical dashed square) and the source-drain current at this operating point (I_DS_ at V_G_ = -0.5 V) was used to construct calibration curves and extract sensor slopes and limits of detection.

## Results

For the specific quantification of ESKAPE pathogens, six DNA aptamers previously published for different applications and each reported to bind one member of this group of bacteria with high affinity, were selected. Their sequences, original literature sources and the key concepts of their discovery are summarized in Table 1 also stating the fact that all aptamers originate from independent researchers and have been selected from literature. They have previously been validated in diagnostic or biosensing contexts, making them promising candidates for integration into the rGO-based FET platform. To evaluate whether these aptamers retain their target specificity on a graphene electronic transducer, each aptamer was immobilized on an individual rGO-FET device. The resulting aptamer-functionalized chips were subsequently exposed to either their corresponding target bacterium (“specific”) or to a mixture containing all remaining ESKAPE members (“non-specific”) across a concentration range of 10^1^-10^5^ CFU mL^-1^. The binding events were monitored via changes in the source-drain current (ΔI_DS_) at a fixed gate voltage of -0.4 V, derived from the respective I_DS_V_G_ curves.

**Table 1:**
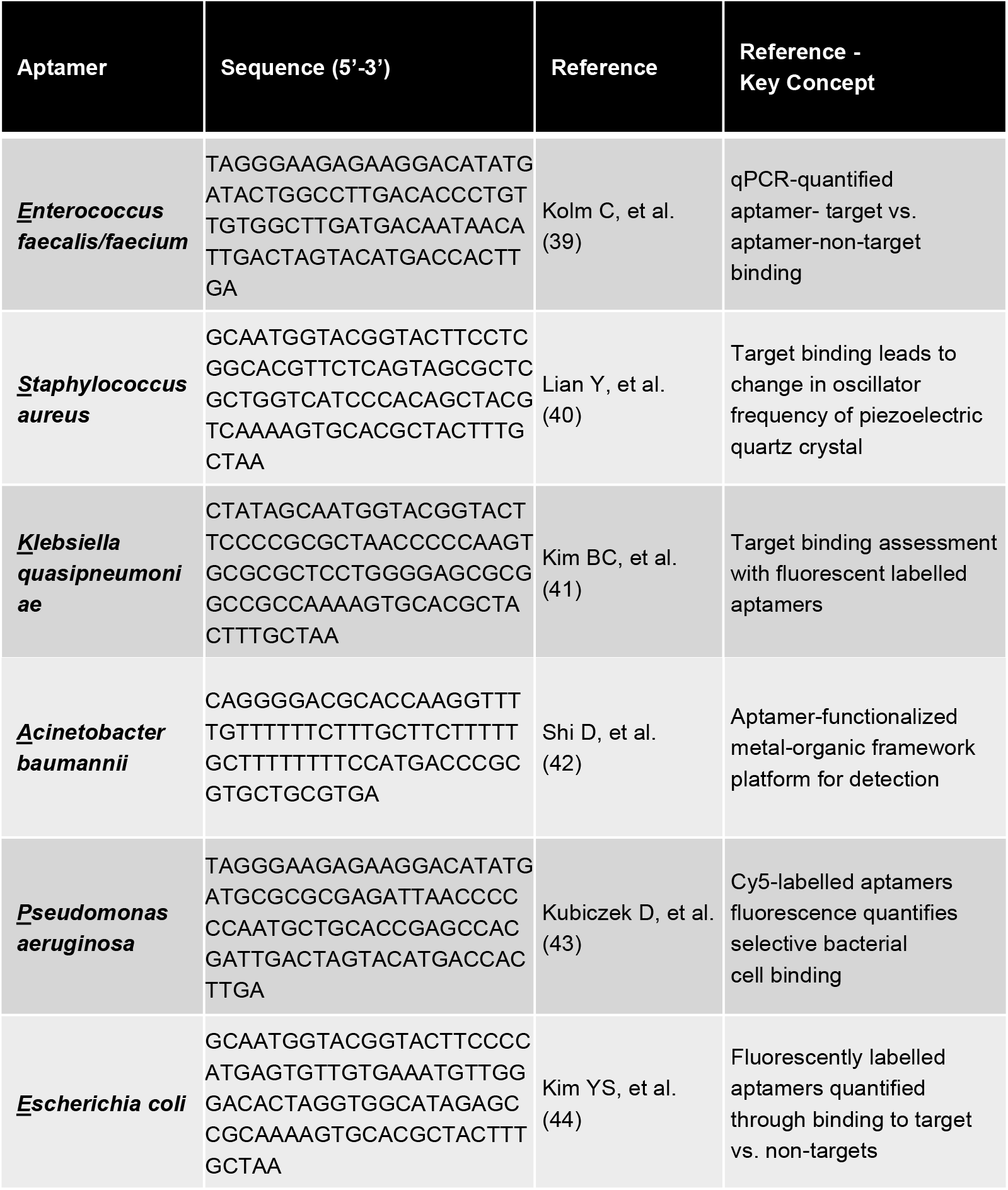
Overview of ESKAPE pathogen specific aptamers collected from the literature, including their sequences, original references and a short key concept describing the core finding of each study.

All aptamer-target pairs exhibited a clear and concentration-dependent increase in ΔI_DS_, demonstrating robust charge modulation induced by specific bacterial binding (Fig 2A). For subsequent quantitative analysis, a fixed gate voltage of -0.4 V was chosen and the source-drain current I_DS_ at this operating point (I_DS_ at V_G_ = -0.4 V) was used to construct calibration curves (Fig S1) and extract sensor slopes and limits of detection (Fig 2B). These signals increase not only provide direct evidence that the selected aptamers maintain their molecular recognition capability when immobilized on rGO and that their interaction with the corresponding pathogen measurably perturbs the carrier density of the rGO channel but also proved that the incubation with non-target bacterial mixtures failed to deliver specific signals (Fig S1, Fig 2A). These irregular low-level fluctuations most likely arise from transient nonspecific adsorption or weak electrostatic perturbations that do not result in stable carrier modulation. The strong contrast between the two conditions (specific, non-specific) highlights both the intrinsic specificity of the aptamers and the ability of the rGO-FET sensor to discriminate true binding events from nonspecific interactions.

**Fig 2:**
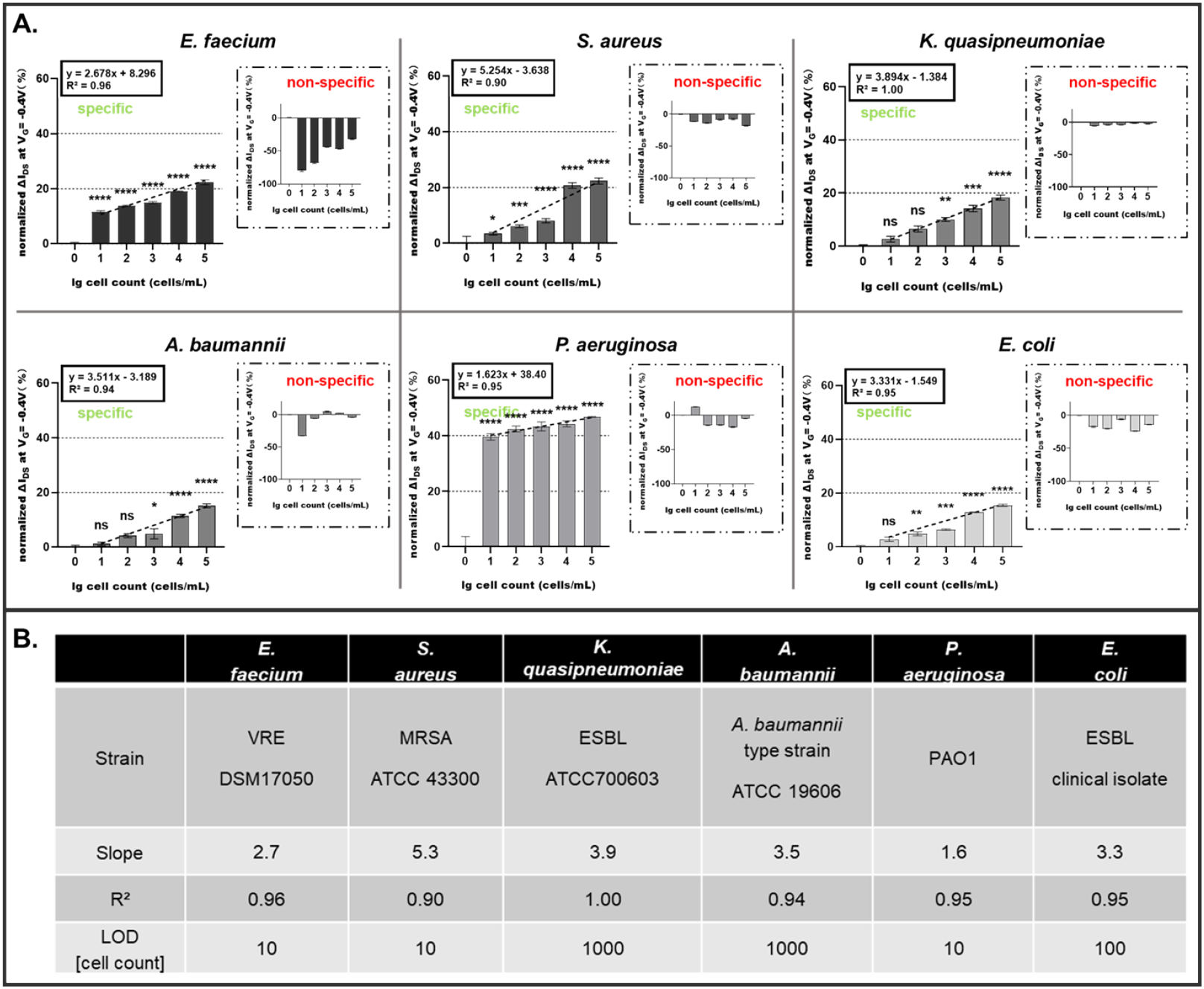
Specificity and analytical performance of aptamer-functionalized gFETs toward ESKAPE pathogens. (A) gFET responses of each aptamer toward its specific (target strain of specific aptamer) and non-specific bacterial targets (mix of all other ESKAPE strains without the specific strain), shown as concentration-dependent signal changes. Specific interactions exhibit positive signal trends, while non-specific interactions remain negative. Measurements were performed with a gate voltage sweep from -0.5 V to +0.5 V; for each measurement point on the chip, three consecutive sweeps were conducted. (B) Summarizes the extracted calibration parameters (slope, R^2^) and the calculated limits of detection (LOD) for each aptamer-target pair. Statistical significance was assessed using a one-way ANOVA performed in GraphPad Prism. A *p*-value < 0.05 was considered statistically significant, where *\** denotes < 0.05, *\*\** denotes < 0.01, *\*\*\** denotes < 0.001 and *\*\*\*\** denotes < 0.0001.

For all pathogens, the specific ΔI_DS_ values obtained across the concentration range exhibited excellent linear correlation with the logarithmic bacterial count, with R^2^ between 0.90 and 1.00 (Fig 2B). The slopes of the linear fits varied between aptamers, reflecting differences in affinity, cell-surface charge signatures and the strength of graphene carrier perturbations. The steepest response was observed for the *S. aureus* aptamer (slope = 5.3), whereas the *P. aeruginosa* aptamer produced a shallower but still robust response (slope = 1.6). Limits of detection calculated from first significant cell count to the zero-cell reference and ranged from 10 to 1000 cells. These results demonstrate that the platform is suited for highly sensitive quantification of bacterial load over several orders of magnitude. Across all six pathogens, the absence of systematic positive signals in the nonspecific controls (all ESKAPEs excluding aptamers target strain) the specific confirms that the aptamers retain high selectivity toward their cognate bacterial surfaces and that the electrical readout faithfully amplifies these binding events without introducing artefactual responses. Collectively, these findings demonstrate that aptamer-functionalized rGO-FETs can resolve specific molecular recognition events with high sensitivity, precision. The consistent performance across all tested ESKAPE pathogens establishes this biosensing architecture as a robust and scalable approach for rapid, label-free and selective electrical detection of clinically relevant multidrug-resistant bacteria.

## Discussion

Three decades after the first reports of nucleic-acid ligands capable of selective molecular recognition, aptamers have become a mature and broadly adopted class of affinity reagents (45)(46). Their evolution through iterative enrichment and diversification strategies has generated libraries of DNA and RNA molecules with binding characteristics comparable to antibodies, while offering advantages in chemical stability, reproducibility and surface immobilization. These properties have supported their integration into a wide range of diagnostic and biotechnological applications. In parallel, electrolyte-gated field-effect transistors (EG-FETs) have developed into an attractive and sensitive bioanalytical toolset (47)(48)(49). These devices respond to local electrostatic changes at the semiconductor-electrolyte interface, allowing biomolecular interactions occurring near the transistor surface to be converted directly into measurable I_DS_ shifts (50). This label-free, rapid mechanism enables compact sensor designs suited for near-patient analysis (51).

Our previous work demonstrated that aptamer-functionalized graphene rGO-FETs can detect a variety of biological targets, including commensal microbiome species and viral pathogens, with high sensitivity and specificity (31)(32)(33)(34)(35). Building on this foundation, the present study expands the platform to the clinically critical ESKAPE pathogens by incorporating six independently derived SELEX aptamers, each validated in different assay formats, to assess the system’s adaptability. All aptamers maintained their specificity after coupling rGO, while cognate pathogens induced clear, concentration-dependent ΔI_DS_ responses, non-target mixture produced only minimal background fluctuations. The limits of detection between 10 and 1000 cells align with the requirements for rapid pathogen diagnostics. This successful use of six literature-derived aptamers highlights the potential of the rGO-FET aptasensors as rapid, selective, scalable diagnostic tool that can be adapted to a broad range of different organisms. Thus, we dare to hope that we have added another piece of evidence for the ease and functionality of rGO-FET-based aptasensors to the portfolio of specific and sensitive quantification tools for diagnostics of microbial pathogens and hope to inspire other researchers to make use of already existing aptamers.

## Material and Methods

### Cell lines and culture conditions

All bacterial isolates were kept on sheep blood agar plates (Oxoid, Basingstoke, UK) for routine maintenance. For liquid growth, *Escherichia coli, Acinetobacter baumannii* and *Pseudomonas aeruginosa* were cultivated overnight in LB-Miller medium at 37°C with orbital shaking at 160 rpm. The other species were propagated in Todd-Hewitt broth (Oxoid) enriched with 0.5 % yeast extract (BD, USA). Unless indicated otherwise, both broth cultures and agar plates were incubated at 37°C. All bacterial strains used in this study were provided by the Institute of Medical Microbiology and Hygiene at the University Hospital Ulm. The panel included *Enterococcus faecium* (DSM 17050), *Staphylococcus aureus* (ATCC 43300), *Klebsiella pneumoniae* (ATCC 700603), *Acinetobacter baumannii* (ATCC 19696), *Pseudomonas aeruginosa* PAO1 and *Escherichia coli* (BSU1286).

### Synthesis and Preparation of Aptamers

The 5’-amino (-NH_2_)-modified aptamers targeting *Enterococcus faecium, Staphylococcus aureus, Klebsiella pneumoniae, Acinetobacter baumannii, Pseudomonas aeruginosa, Escherichia coli* were custom synthesized by biomers (Ulm, Germany). Before use, the aptamers were dissolved in 1× DPBS to a final concentration of 100 pmol.

### Surface Modification of rGO-FET for Biosensing

Field-effect transistors with reduced graphene oxide (rGO) as the channel material (rGO-FETs) were fabricated following the procedure reported by Kissmann et al. (52). Surface functionalization was performed by incubating the rGO-FET in a dimethyl sulfoxide (DMSO) solution containing 500 mM Hydroxypyrene and 50 μM 1-pyrene carboxylic acid (PCA) at room temperature for 12 h. The device was then rinsed with isopropanol, dried under nitrogen flow and activated with a 0.01× DPBS (phosphate-buffered saline) solution containing 15 mM 1-ethyl-3-(3-dimethylaminopropyl) carbodiimide (EDC) and 15 mM N-hydroxysuccinimide (NHS) for 30 min to promote carboxyl group activation. Subsequently, 100 nM of 5’-amino (-NH_2_)-modified aptamers in 1× DPBS were covalently immobilized onto the surface. After a final rinse with 0.01× DPBS to remove unbound aptamers, the biosensor was ready for electrical detection of ESKAPE group.

### Electrical Detection of ESKAPE Using rGO-FETs

ΔI_DS_ curves were recorded using a KEYSIGHT Modular USB Source Measure Unit by sweeping V_G_ from –0.5 to 0.5 V. Higher voltages were avoided to prevent damage to the bio-functional components. Bacteria of ESKAPE in logarithmic growth with OD=0.8 were diluted stepwise to obtain different numbers of bacteria from 0 to 10^5^. The microfluidic channel was first flushed with bacterial suspension at 0.5 mL/min for 1 min to promote adhesion, followed by a 15 minute equilibration at 0.2 mL/min to mimic physiological flow, and finally washed with 1% DPBS at 0.5 mL/min for 5 min to remove unbound cells, ensuring optimal conditions for FET-based biosensing.

## Supporting information

**Figure S1:**
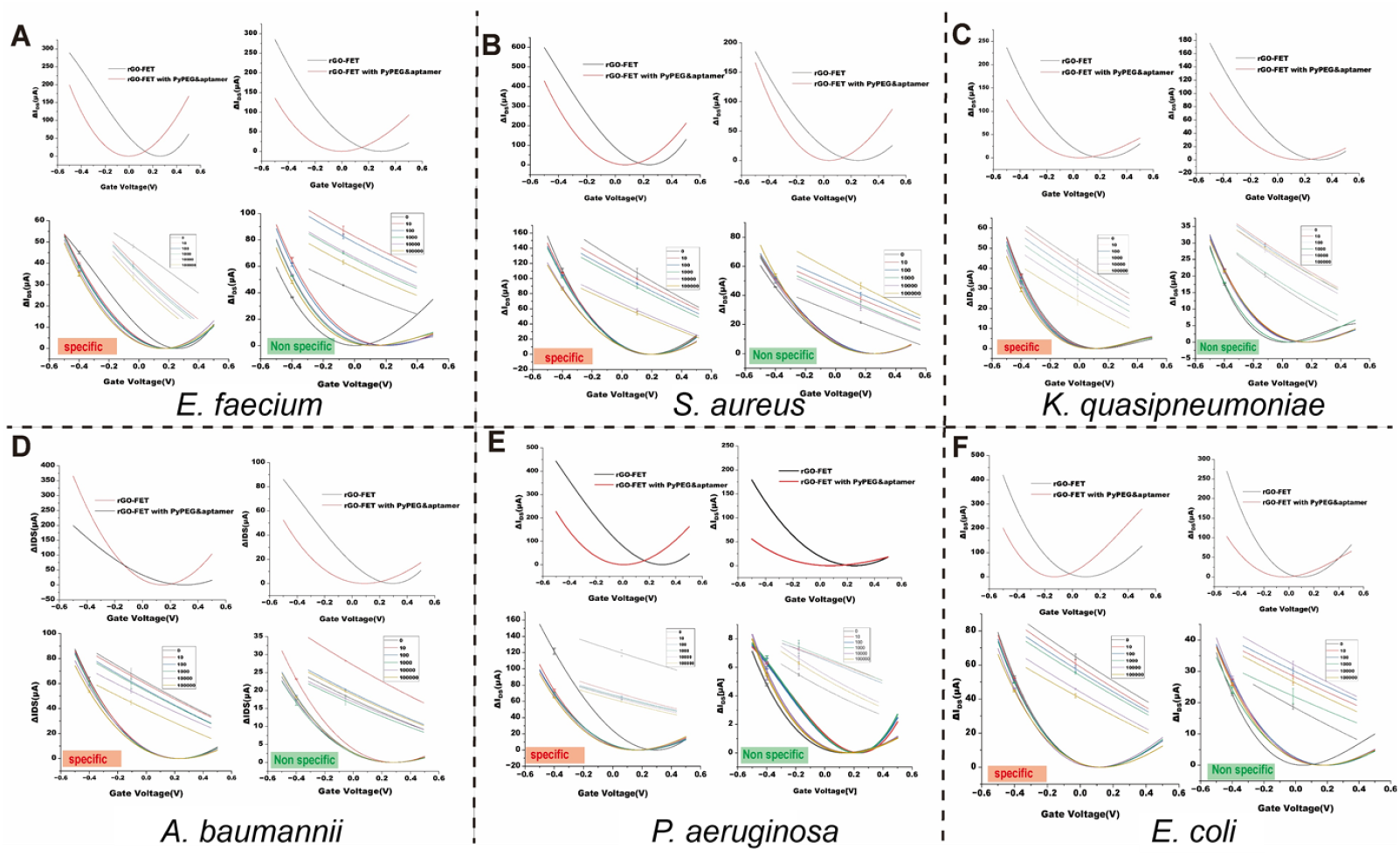
Transfer characteristics (I_DS_V_G_ curves) of aptamer-functionalized rGO-FET device exposed to their corresponding ESKAPE pathogen (specific) or a non-target bacterial mixture. Each panel (A-F) displays the electrical response for devices functionalized with aptamers against (A) E. faecium, (B) S. aureus, (C) K. quasipneumoniae, (D) A. baumannii, (E) P. aeruginosa and (F) E. coli. For each condition diluted bacterial suspensions (10^1^ -10^2^ CFU/mL) were introduced into the microfluidic system and the resulting device response were recorded as changes in source-drain curren with a gate voltage sweep from -0.5 V to +0.5 V, For each measurement on the chip, three consecutive sweeps were conducted.

## Data availability statement

Data available at: https://data.mendeley.com/datasets/wrkrs83rm8/1

